# Screening for metastasis-related genes in mouse melanoma cells through sequential tail vein injection

**DOI:** 10.1101/2024.01.29.577709

**Authors:** Qinggang Hao, Junling Shen, Lei Sang, Yan Bai, Jianwei Sun

**Affiliations:** Center for Life Sciences, Yunnan Key Laboratory of Cell Metabolism and Diseases, State Key Laboratory for Conservation and Utilization of Bio-Resources in Yunnan, School of Life Sciences, Yunnan University, Kunming 650500, China

**Author notes:** Correspondence to: Jianwei Sun.

**Keywords:** Melanoma, Lung metastasis model, Gene screening, Tail vein injection

## Abstract

Tumor metastasis, responsible for approximately 90% of cancer-associated mortality, remains poorly understood. Here in this study we employed a melanoma lung metastasis model to screen for metastasis-related genes. By sequential tail vein injection of mouse melanoma B16F10 cells and the subsequently derived cells from lung metastasis into BALB/c mice, we successfully obtained highly metastatic B16F15 cells after five rounds of *in vivo* screening. RNA-sequencing analysis of B16F15 and B16F10 cells revealed a number of differentially expressed genes, some of these genes have previously been associated with tumor metastasis while others are novel discoveries. The identification of these metastasis-related genes not only improve our understanding of the metastasis mechanisms, but also provide potential diagnostic biomarkers and therapeutic targets for metastatic melanoma.

## 1. Introduction

Cancer has emerged as the second leading cause of death world-wide, posing a significant challenge to public health. Despite the rapid advancement in medicine and the ongoing improvement in treatment methods, tumor recurrence and metastasis persist as crucial factors contributing to poor prognosis, morbidity and death in cancer patients (Christofori 2006). More than 90% of cancer -related death are attribute to distant metastasis (Chaffer and Weinberg 2011). Tumor metastasis, a complex multi-step process, hinges on the characteristic changes in tumor cell phenotypes, accompanied by genetic and epigenetic alterations in the tumor cell. However, the genes that governing tumor metastasis, especially in melanoma, largely remain elusive (Turajlic and Swanton 2016). Therefore, exploring new and effective factors that regulate tumor metastasis is urgently needed.

Tumor heterogeneity, defined by the presence of diverse cell populations within a single tumor, contribute to variations in cell proliferation rates, morphology, genetic makeup, surface markers, metabolism, and the ability to infiltrate tissues, metastasize, and respond to treatments. A challenge in treating heterogenous tumors arise from the different responds of various cell types. Lower differentiation in a tumor, indicative of tumor cells deviating from normal cells, often correlates with increased heterogeneity and suggests a more aggressive cancer types that may be more challenging to treat effectively and more prone to metastasis. Hence, reducing tumor heterogeneity and isolating highly metastatic cells are crucial steps in identifying key genetic factors that drive metastasis. This approach may lead to the discovery of potential therapeutic particularly in cancers where metastasis significantly worsens the prognosis (Bai *et al*. 2023; Fidler 1973; Minn *et al*. 2005). Understanding the genetic basis of tumor metastasis is vital for developing treatments that can more effectively combat these highly aggressive and often fatal cancer types.

In this study, we performed sequential tail vein injection of B16F10 and subsequent derived cells from lung metastases (Yang *et al*. 2012), successfully obtaining highly metastatic B16F15 cells. RNA-seq analysis of B16F10 and B16F15 cells revealed various metastasis genes. This approach not only contributes to the scientific understanding of the genetic factors driving metastasis, but also holds practical implications for developing new treatments for metastatic melanoma.

## 2. Experimental Section

### 2.1 Materials and Reagents

1. RPMI 1640 and DMEM medium were supplemented with 10% FBS and and 1% penicillin-streptomycin.
2. 0.25% trypsin (add 1mM EDTA), 1 mg/mL type IV collagenase.
3. 100 mg/mL kanamycin and 100 mg/mL ampicillin.
4. 10cm cell and tissue culture dishes.
5. 1× PBS (Phosphate buffered saline).
6. 10mg/mL D-luciferin.
7. Dissecting scissors, scalpel and forceps.
8. Tanon ABL-X5 small animal live imaging system.
9. 4-6 week old BALB/c mice were obtained from the Animal Research and Resource Center, Yunnan University. {Certification NO. SCXK(Dian)K2021-0001}. All animal work procedures were approved by the Animal Care Committee of the Yunnan University (Kunming, China).
10. B16F10 and 293T cell lines.

### 2.2 B16F10 cells culture and treatment

1. The B16F10 cells were cultured in RPMI 1640 medium supplemented with 10% FBS in a 5% CO_2_ culture incubator at 37 °C.
2. When the cells reached to 80-90% confluence, the medium was discarded, and cells were gently washed twice with PBS. Subsequently, cells were harvested by trypsinization.
3. After approximately one minute, when the cells detached from the culture dishes, trypsinization was halted by adding culture medium. Cells were then collected into the centrifuge tube.
4. The collected cells were centrifuged at 2000 rpm for 3 minutes at room temperature, the supernatant was discarded and the cells were resuspended in 1 mL medium. Cell number was determined using a hemocytometer, and cells were diluted with RPMI 1640 medium.

### 2.3 Establishment of luciferase labeled B16F10 stable cell line by lentiviral infection

1. Cultured 293T cells in DMEM medium supplemented with 10% FBS.
2. Approximately 12 hours before transfection, seeded the 293T cells onto a 10cm cell culture dish. Ensure cells reached 80–90% confluence and changed medium approximately 2 hours before transfection.
3. Transfection and viral packaging. Mixed 5 μg pLenti PGK Blast V5-LUC (w528-1) (plasmid# 19166), 5 μg each of the packaging plasmids VSV-G (plasmid# #138479) and psPAX2 (plasmid #12260) in 0.5 mL of serum-free medium, mixed 15 μg PEI into 0.5 mL of serum-free medium, vortexed for one minute, and then incubated for 5 minutes. Mixed the plasmids with the PEI solution, vortexed for one minute, and incubated the mixture for 15 min. Dropwise, added the mixture into the 10cm 293T cell culture dish, and changed fresh medium 24 hours after transfection.
4. Harvested lentivirus from the medium 48 hours and 72 hours after transfection. Filtered the lentivirus solution through a 22 μm filter.
5. Added 10 mL of the filtered lentivirus to infect B16F10 cell. Feeded the cells with fresh medium after 48 hours (serum can be added halfway). Screened successfully infected cells using blasticidin and measured the luciferase activity of B16F10 cellsbefore tailvein injection.

### 2.4 Tail vein injection of luciferase labeled B16F10 cells into BALB/c mice

1. When B16F10 cells reached to 80-90% confluence, harvesedt the cells by trypsin digestion, determined the cells with a hemocytometer, and diluted the cells with PBS to a concentration of 1×10^8^ cells/mL.
2. Dilated the caudal vein of mice using a heating lamp. Fix the mouse in a fixator and expose its tail..
3. Mixed the B16F10 cells and load 120 μL into the syringe fitted with a 27G needle. Keep the needle bevel level with the syringe scale.
4. Removed the bubbles from the syringe and needle, and injected B16F10 cells into one of the lateral tail veins.
5. Ensure successful injection at least three mice.

### 2.5 Bioluminescence imaging and analysis

1. Intraperitoneal injection of 100 μL 10 mg/mL D-luciferin per mouse.
2. Anesthetized mice with isoflurane 5 minutes later.
3. Subjected the mice to live imaging, recorded the luminescence signal with Tanon ABL-X5 small animal live imaging system. Captured images within 1 hour of the first day and weekly after tail vein injection.

### 2.6 Screening of high metastatic melanoma cells

1. The mice were sacrificed when the luminescence signal was detected through Tanon ABL-X5 small animal live imaging. Disinfected the sacrificed mice in 75% alcohol for 20 seconds. Dissected and obtained the lungs in a clean hood.
2. The lungs were washed with PBS for twice, then transfer to 10cm cell culture dish, chop the lung into pieces as small as possible with sterilized scalpel and then add 2 mL 1 mg/mL collagenase IV.
3. Added 4 mL RPMI 1640 medium, mix well and digest the lung tissue at 37^0^C for 30 minutes.
4. Removed collagenase by centrifuging and discarded the supernatant. Cultured the lung tissue and metastatic melanoma cell in RPMI 1640 medium supplemented with 5 ug/mL blasticidin, 100 ug/mL kanamycin and 100 ug/mL ampicillin. (Blasticidin is used to screen for metastatic tumor cells, while kanamycin and ampicillin are used to prevent bacterial contamination).
5. The cells obtained through this round of screening are called B16F11, the cells were then harvested for the next round of screening. After 5 rounds of screening, B16F15 cells were obtained.

### 2.7 RNA-seq

1. Cultivated B16F10 and B16F15 cells to a comparable state, collecting 1×10^7^ cells in each culture dish with Trizol Reagent. Ensure three biological replicates for each sample.
2. Library construction and RNA-seq were performed by <u>Novogene</u> company using an Illumina NovaSeq 6000 platform. The sequencing quality was assessed using FastQC.
3. The index construction is performed using bowtie2 (http://bowtie-bio.sourceforge.net/bowtie2/index.shtml), the sequence alignment of the clean data is conducted using TopHat2 (https://ccb.jhu.edu/software/tophat/index.shtml), and the transcriptome assembly is performed using cufflinks (https://cole-trapnell-lab.github.io/cufflinks/).
4. The expression level of each gene was calculated according to the fragments per kilobase million reads (FPKM) method.
5. The differential gene expression analysis is performed using DEG seq2 log_2_(B16F15 mean (FPKM)/B16F10 mean (FPKM)). The p-value is calculated using the t-test. Different expressed genes (DEGs) with |log_2_(B16F15/B16F10)| ≥ 1, and P-value < 0.05 were considered to be significant (DEGs).
6. The DEGs were used to perform pathway analysis including Gene Ontology (http://www.geneontology.org) and Kyoto Encyclopedia of Genes and Genomes (http://www.genome.jp/7eg/) by using cluster profiles with adjusted P-value ≤0.05.

### 2.8 Statistical analysis

GraphPad Prism 9 were used for the basic statistical analysis.

## 3. Results and Discussion

### 3.1 The cells derived from the sequential tail vein injection lung metastasis model possess enhanced metastatic ability

To identify melanoma metastasis-related genes, B16F10 cells stably expressing luciferase were generated. Subsequently 1×10^6^ B16F10 cells were injected into the BALB/c mice via the tail vein. Upon detecting a significant fluorescence luminescence signal in the lungs through *in vivo* imaging, the mouse was sacrificed and the cells were collected and selected using blasticidin. The cells termed B16F11 were cultured for the next round of tail vein injection. Highly metastatic B16F15 cells were obtained after 5 rounds of *in vivo* and *in vitro* screening, demonstrating significantly enhanced metastatic activity of melanoma cells from each careening round (**Figure 1A**).

**Figure 1.**
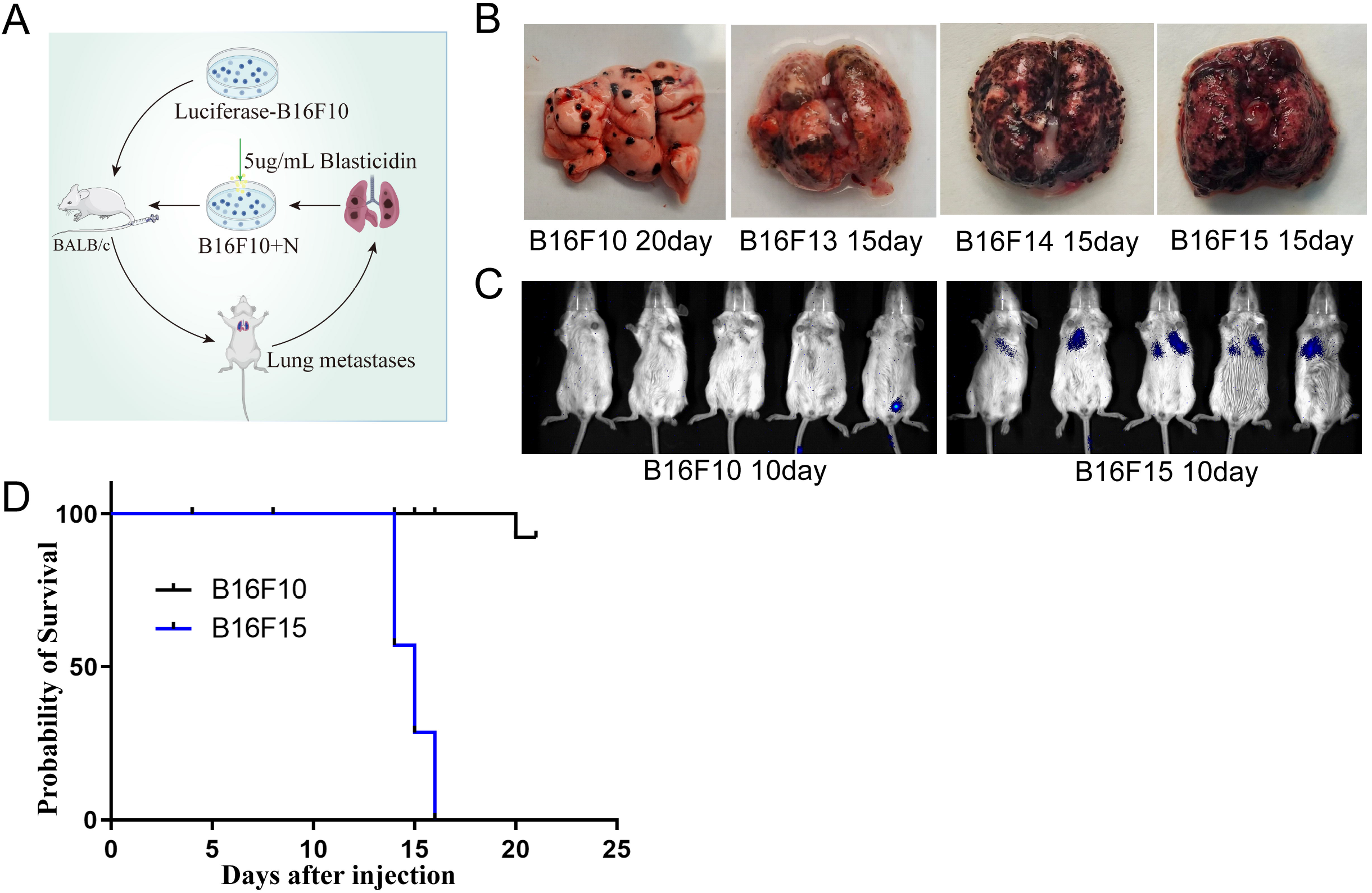
Screening of highly metastatic melanoma cells. (A) Scheme of highly metastatic tumor cell screening. (B) Representative images of lung metastasis in different rounds of *in vivo* screening. (C) Representative images of live animal imaging to assess lung metastasis after tail vein injection of B16F10 and B16F15 cells in BALB/c mice. (D) Survival analysis of BALB/c mice after tail vein injection of B16F10 and B16F15 cells.

To validate the screening, we injected B16F10 and B16F15 cells into mice via the tail vein respectively, confirming the markedly stronger metastatic activity of B16F15 compared to the original B16F10 cells (**Figure 1B and 1C**). Survival analysis of the mice bearing lung metastases revealed substantially shorter survival time for B16F15 mice compared to B16F10 (**Figure 1D**).

### 3.2 Identification of differentially expressed genes (DEGs) and thieir role in metastasis pathways

To uncover key genes that may be involved in regulating tumor metastasis, B16F10 and B16F15 cells were collected for RNA-seq analysis. A substantial number of DEGs were identified (|Log_2_FC| >= 1, P value < 0.05). Among them, 1329 DEGs were significantly up-regulated, and 1581 were remarkably down-regulated in B16F15 cells (**Figure 2A-2B**). Several well-known metastasis genes were included in the list of DEGs, including TCF7 (Zhao *et al*. 2023), MAPK13 (Tan *et al*. 2010), RAB17 (Zhou *et al*. 2021), CXCL10 (Ka *et al*. 2023; Lee *et al*. 2022; Shang *et al*. 2022), CTSK (Khan *et al*. 2021; Li *et al*. 2019; Wu *et al*. 2022), CCDC80 (Huang *et al*. 2023), and ABCC2 (Sensorn *et al*. 2016). In line with RNA-seq analysis, our qRT-PCR confirmed the elevated expression of the aforementioned DEGs (**Figure 2C**). All these findings demonstrate the feasibility of our approach.

**Figure 2.**
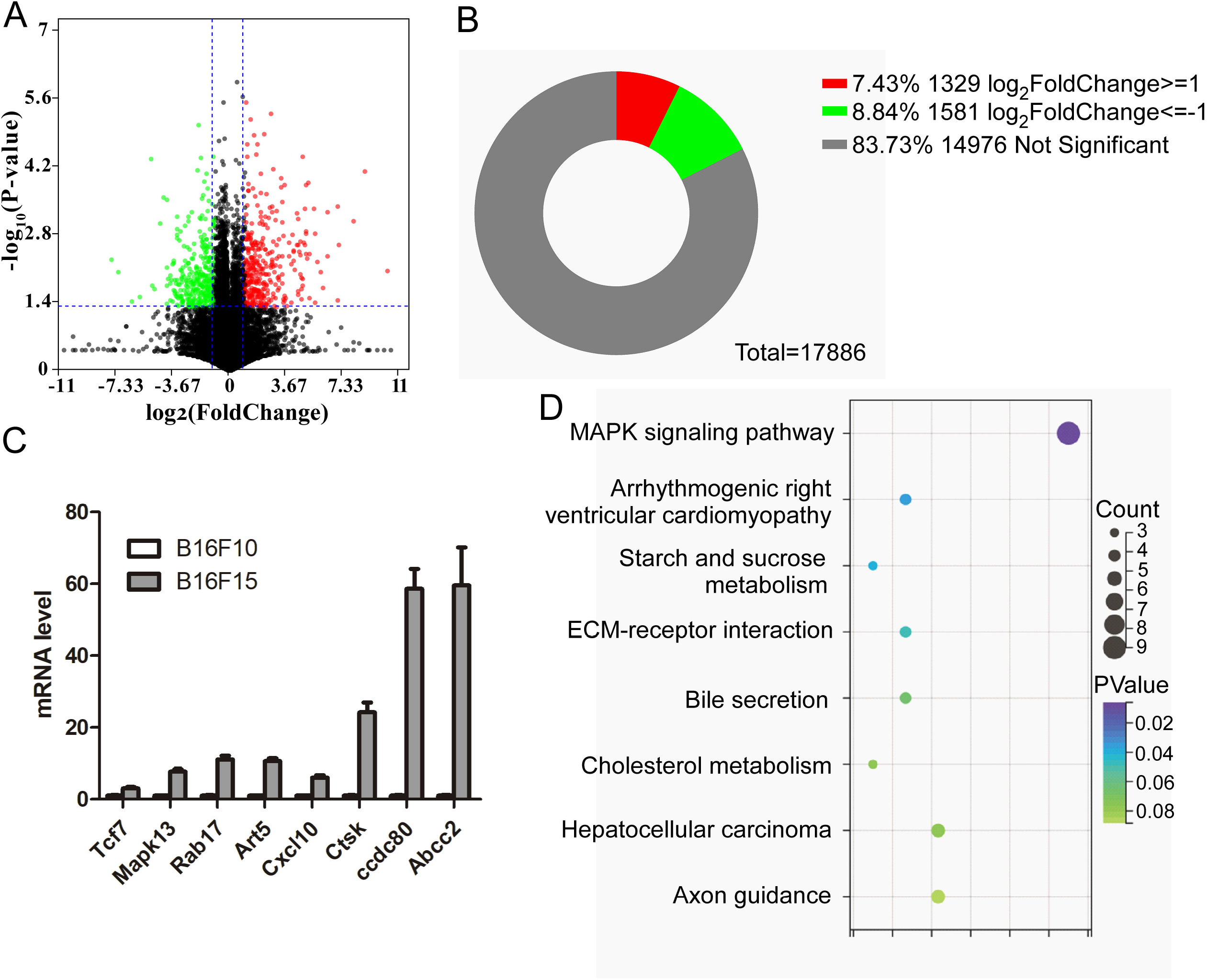
Identification of DEGs between B16F10 and B16F15 cells. (A) Volcano plot analysis of DEGs between B16F10 and B16F15. (B) Statistical analysis of the number of DEGs. (C) qRT-PCR analysis of expression levels of indicated genes in B16F10 and B16F15 cells. (D) The enrichment analysis of DEGs between B16F10 and B16F15 cells.

To investigate the function of these DEGs, we performed KEGG analysis and found that these DEGs were involved in ECM, the MAPK pathway, glucose and cholesterol metabolism (**Figure 2D**). These findings may provide potential diagnostic biomarkers and therapeutic targets for metastatic melanoma.

### 3.3 Discussion

The tumor metastatic cascade is a complex series of steps that cancer cells undergo to spread from the primary tumor to distant sites in the body. The steps include: 1. Epithelial-mesenchymal transition (EMT): this process allows epithelial cells, which are normally stationary, to acquire the ability to move and behave more like mesenchymal cells, which are more fibrous and mobile; 2. Invasion: the cells then invade tissues nearby, including entry into blood vessels (intravasation) or lymphatic systems; 3. Circulation: the cells travel through the circulatory system to distant sites; 4.Extravasation: the cells exit the circulatory system, which involves interacting with the microvessels of distant tissue; 5. Colonization: finally, the cells establish new tumors in these distant sites.

The tail vein injection model of lung metastasis is a well-established experimental system for studying these processes *in vivo* (Fidler 1973; Minn *et al*. 2005). In this study, we injected luciferase-labeled melanoma B16F10 cells into the tail vein, after which they circulate and establish lung metastases. Luciferase allows us to track the cells using *in vivo* luminescence imaging, which is a powerful technique for visualizing the location and growth of metastatic tumors in live animals. This model is valuable for identifying genes and signaling pathways involved in metastasis by comparing the primary tumor cells with those that have successfully metastasized. Recent studies have utilized genome-wide microarrays to compare gene expression profiles across different stages and models of metastasis. Specifically, significant differences were observed between primary orthotopically implanted tumors and their lung metastases, but not between lung metastases arising from orthotopic implantation and those from tail vein injections. This suggests that despite the method of metastasis induction, once cancer cells reach the lungs, the genes they express may converge to facilitate survival and growth in this new environment. This information is invaluable because it supports the use of experimental metastasis models to identify genes involved in the metastatic process. It also indicates that cells with a high metastatic potential may pre-exist within the heterogeneous primary tumor population. The tumor microenvironment and the selective pressures it imposes could be key factors in determining which cells ultimately metastasize.

The candidate genes identified in our study are hypothesized to play roles in: 1. Tumor cell stemness: this refers to the ability of some cancer cells to exhibit stem cell-like properties, which can lead to tumor growth and resistance to conventional therapies; 2. Immune escape: the ability of tumor cells to evade detection and destruction by the immune system, which is a critical hurdle in the fight against cancer; 3. Promotion of metastasis: facilitating the spread of cancer cells from the primary site to distant organs. Further validation of these candidate genes can clarify their roles in the metastatic process and potentially reveal new targets for therapeutic intervention. Understanding these mechanisms can lead to the development of strategies to prevent the spread of tumors, which is crucial for improving outcomes in cancer treatment.

## 4. Conclusion

In conclusion, we have identified DEGs through sequential tail vein injection of the mouse melanoma metastasis model. Some of the genes have been previously reported to be involved in tumor metastasis, while others are novel. Our future work will aim to elucidate the role of these DEGs in melanoma metastasis.

## ACKNOWLEDGMENT

We thank Dr. Jing Li for critical reading of the manuscript. This work was supported by the National Natural Science Foundation of China (82273460 and 32260167), Yunnan Applied and Basic Research Program (202101AV070002), Major Science and Technique Programs in Yunnan Province (202202AA310207), Yan’an Hospital Affliated to Kunming Medical University, grants (KC-23233927, KC-23234451, ZC-23236399, ZC-23236296 and TM-23236924) from Yunnan University.

## Compliance with Ethical Standards

### Conflict of interest

Qinggang Hao, Junling Shen, Lei Sang, Yan Bai and Jianwei Sun declare that they have no conflict of interest.

### Animal rights and informed consent

All institutional and national guidelines for the care and use of laboratory animals were followed.

## Notes

### Competing Interest Statement

The authors have declared no competing interest.

